# The molecular determinants of PABPC-mediated deadenylation rate

**DOI:** 10.64898/2026.06.05.728831

**Authors:** Ryan Y Muller, Tanner M Myers, Eugene Valkov, David P Bartel

## Abstract

Deadenylation, the enzymatic shortening of the poly(A) tail, is typically the first committed step of mRNA decay. Deadenylation rates span nearly a 1000-fold range between transcripts and are governed by protein–RNA interactions, including those involving cytoplasmic poly(A)-binding protein (PABPC). Previous work shows that PABPC can straddle the junction between the poly(A) tail and the 3′ untranslated region (UTR), but whether this conformation influences deadenylation has not been tested. To investigate how straddling influences deadenylation kinetics, we designed a library of tailed RNA substrates and measured both in vitro deadenylation rates in the presence of PABPC and PABPC binding propensity for each substrate. We found that 3′ UTR sequences influence deadenylation through two mechanisms. First, structured UTRs are deadenylated more slowly in the absence of PABPC1, an effect that is alleviated by PABPC1. Second, sequences upstream of the poly(A) tail modulate PABPC1 binding propensity, with tighter binding correlating with slower deadenylation. This relationship is abolished with a PABPC1 mutant lacking UTR-binding capacity. Together, these results show that sequences upstream of the poly(A) tail tune PABPC1 binding and deadenylation rates, likely contributing to the range of deadenylation rates observed for cellular mRNAs.

## Introduction

Messenger RNA (mRNA) decay is a central determinant of gene expression, shaping both the level and timing of protein production. In eukaryotes, most cytoplasmic mRNAs undergo decay through ordered steps starting first with deadenylation—the shortening of the poly(A) tail, then removal of the 5′ cap, and finally 5′-to-3′ exonucleolytic decay^1,2^. Deadenylation is typically both the first committed and the rate-limiting step of mRNA decay^3^.

Regulatory information controlling deadenylation is predominantly encoded in the 3′ untranslated region (UTR), where sequence and structural elements are recognized by microRNAs and RNA-binding proteins (RBPs). MicroRNAs recruit silencing complexes that engage deadenylases to accelerate poly(A)-tail shortening, while many RBPs bind defined 3′ UTR elements and recruit deadenylation machinery to accelerate poly(A)-tail shortening^4–8^, for example, RBPs such as Pumilio, Tristetraprolin (TTP), Roquin, and AUF1 each bind specific elements in the 3′ UTR and recruit deadenylation machinery^9–13^.

Nearly all eukaryotic mRNAs in the cytoplasm carry a 3′ poly(A) tail, which is bound by cytoplasmic poly(A)-binding protein C (PABPC)^14^. PABPC1 is the most abundant and ubiquitously-expressed human paralog, whereas PABPC4 shows lower expression across tissues and PABPC1L is the dominant PABPC paralog expressed in embryonic cells^15,16^. PABPC promotes both translation and mRNA stability, and is displaced from the poly(A) tail during deadenylation^17^.

Through direct binding to the poly(A) tail, as well as recruitment of deadenylation machinery, PABPC1 is capable of both inhibiting and promoting deadenylation^18,19^. PABPC1 binds transcripts along the length of the poly(A) tail using a series of four RNA-recognition motifs (RRM1–4), arranged linearly from the N-terminus. PABPC1 binding to the poly(A) tail provides a steric block from cellular deadenylases, including the CCR4-NOT complex, which is the main deadenylase in humans^20^. At the same time, the C-terminal MLLE domain, also known as the PABPC domain, mediates protein-protein interactions, notably binding to short PAM2 motifs present in a variety of translation and decay factors, such as eIF4G, PAIP2, Tob1/2, and TNRC6^21,22^. Thus, PABPC1 can help to both stabilize and destabilize mRNA, but how the balance between these two processes is determined remains poorly understood.

Each RRM of PABPC1 contributes to RNA binding, but the individual domains differ in both specificity and affinity. Structural and biochemical studies show that RRM1 and RRM2 form a cooperative unit that preferentially binds polyadenosine stretches with high affinity, exhibiting stringent sequence selectivity for homopolymeric A^23–25^. In contrast, RRM3 and RRM4 have relaxed sequence specificity, allowing them to either bind additional poly(A) sequence or interact with the adjacent 3′ UTR^24^. The PABPC1 molecule closest to the 3′ UTR may therefore straddle the poly(A)/UTR junction, with UTR-specific contacts potentially tuning its function on different transcripts.

Here, we investigated how poly(A)-proximal 3′ UTR sequences influence deadenylation kinetics through PABPC1. Sequence adjacent to the poly(A) tail can directly influence deadenylation through its interaction with PABPC1, independent of known regulatory factor binding sites. Thus, PABPC1 protects transcripts from deadenylation not only through poly(A)-tail binding but also via sequence-specific interactions with the adjacent 3′ UTR, providing a mechanism for transcript-specific regulation of mRNA stability.

## Results

### PABPC binding and UTR structure inhibit deadenylation in vitro

Because PABPC1 can interact with the 3′ UTR, we asked whether poly(A)-proximal sequences influence deadenylation and contribute to the nearly 1000-fold range in cellular deadenylation rates^19,26^. To test this model, we designed a 38,700-member library of RNA substrates that mimicked the ends of endogenous transcripts, favoring those transcripts with higher sequence conservation and expression. Each substrate consisted of a 50-nucleotide (nt) segment from the end of a human 3′ UTR followed by a 30-nt poly(A) tail.

To determine how long PABPC1 takes to reach binding equilibrium, we incubated PABPC1 with a mix of radio-labeled A_0_, A_15_, and A_30_ substrate at varying time increments followed by filter binding (Figure S1A). Our expectation was that the A_30_ substrate would have the highest affinity for PABPC1 over A_15_ and A_0_ substrates. Our binding assay required 20 h to see this preference bear out. We did not see significant RNA degradation over this time increment and noted that the abundance of bound RNA scaled directly with the amount of PABPC1 added to the pre-incubation, meaning that PABPC1 retains its ability to bind RNA and also maintains preference for A_30_ over A_15_ and A_0_ over the course of the pre-incubation.

After pre-incubating the library of RNA substrates for 20 h with either excess PABPC1 or no PABPC1, purified CNOT6/7 deadenylase was added (Figure 1A). RNA from replicate reactions was collected at successive time points and sequenced via AVITI 300-nt single-end read to determine tail lengths for each library member (Figure S1B). PABPC1 globally slowed deadenylation (Figure 1B). In its absence, deadenylation was faster and more variable across library members, as indicated by the broader tail-length distributions at later time points (Figure 1B). We hypothesized that intrinsic RNA structure might account for the variation in deadenylation rates observed without PABPC1. Accordingly, we used RNAfold^27^ to predict the minimum free energy of folding (MFE) for each 80-nt sequence (50 nt of UTR with an A_30_ tail) and implemented a linear regression to model deadenylation rates of each library member. We observed a relationship between predicted RNA structure and fitted deadenylation rate, in which RNAs with greater predicted stability were deadenylated more slowly (Figure 1C). Whereas typical deadenylation rates were ∼0.45 nt/min, the most structured substrates were deadenylated at ∼0.2 nt/min or less. These results indicate that intrinsic RNA structure contributes substantially to the variation in deadenylation rates observed without PABPC1.

**Figure 1.**
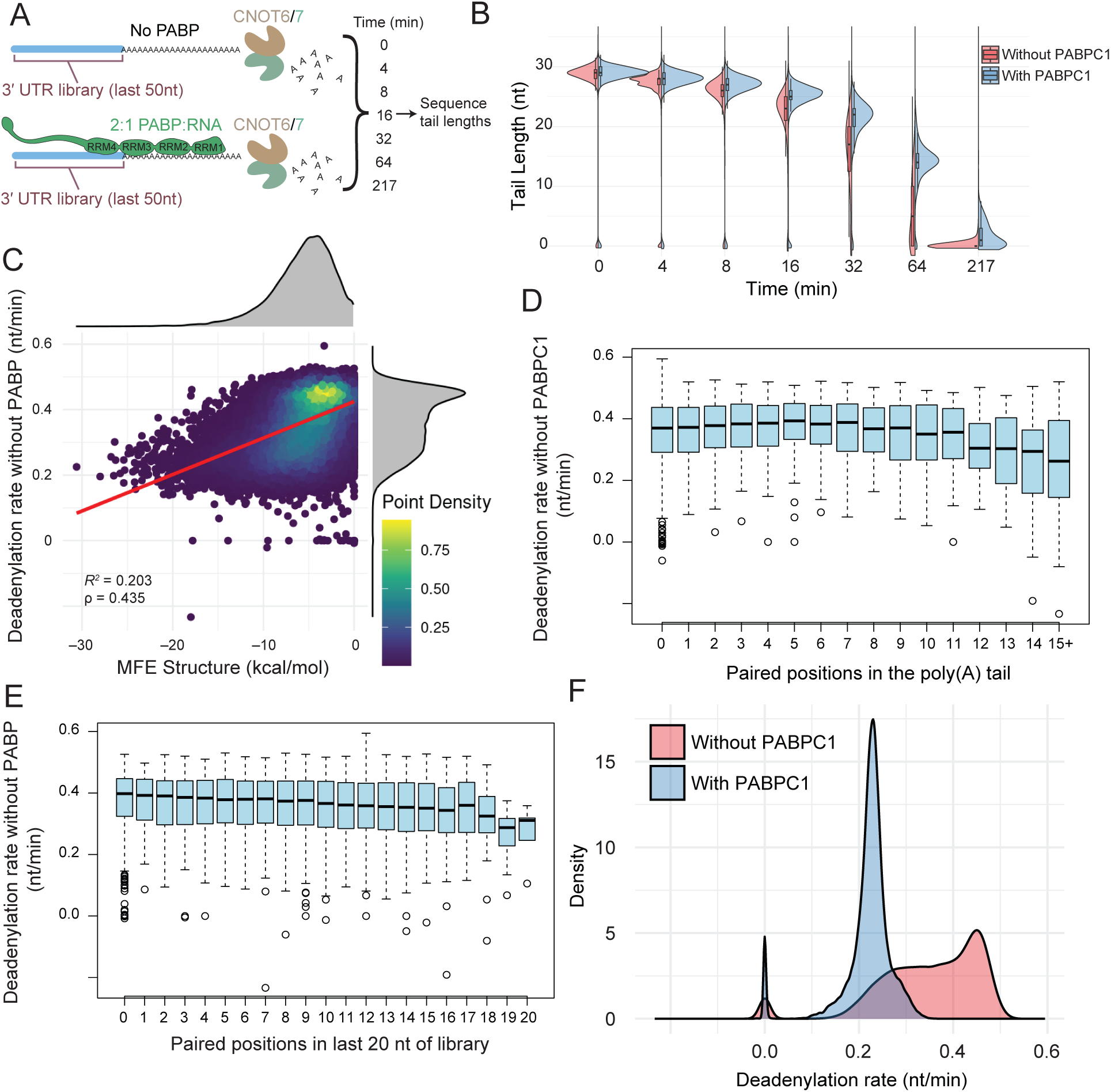
PABPC binding and UTR structure inhibit deadenylation in vitro. (A) Schematic of in vitro deadenylation experiment. A library of poly(A)-tailed RNA substrates is incubated with CNOT6/7 heterodimer in the presence or absence of purified PABPC1. RNA is collected at successive time points, and tail lengths are determined through sequencing. (B) Slower and more uniform deadenylation in the presence of PABPC1. Shown for each time point are distributions of tail lengths in either the absence or presence of PABPC1 (red or blue, respectively). (C) The relationship between in vitro deadenylation and predicted structure. For each member of the library, the fitted deadenylation rate in the absence of PABPC1 is plotted as a function of the minimum free energy (MFE) of folding predicted for the 50 nt of UTR with an A_30_ tail. (D) Reduced deadenylation rates of molecules with more tail residues that are predicted to be paired. Box and whisker plots show the distributions of fitted deadenylation rates in the absence of PABPC1 for library molecules with the indicated number of poly(A)-tail positions that are predicted to be paired in the lowest-energy structure (line, median; box quartile; whisker, 1.5 times the interquartile range (IQR) rule via Tukey’s method). (E) Reduced deadenylation rates of molecules with more tail-proximal residues that are predicted to be paired. This panel is as in D, but instead groups deadenylation rates by the number of positions within 20 nt of the poly(A) tail that are predicted to be paired in the lowest-energy structure. (F) Slower and more uniform deadenylation rates in the presence of PABPC1. Plotted are the distributions of fitted deadenylation rates in either the presence or absence of PABPC1. The peaks at zero are from the A_0_ spike-in controls within the library.

Hypothesizing that structure near or within the poly(A) tail would have a greater effect on deadenylation than more distal structure, we examined how the position of predicted base-pairing affected deadenylation rates. For each A_30_-tailed RNA substrate, the lowest-energy conformation was predicted to determine the number of poly(A)-tail nucleotides predicted to be paired. As the number of poly(A)-tail positions predicted to be paired increased, deadenylation rates tended to slow (Figure 1D). This effect was observed only in the absence of PABPC1; when PABPC1 was added, this structure-dependent deadenylation inhibition was not observed (Figure S1C), suggesting that RNA structure involving the poly(A) tail affects deadenylation only when PABPC1 is absent. We also observed a shallower but monotonic decrease in deadenylation rate with increasing paired positions in the last 20 nt of the variable region, proximal to the poly(A) tail (Figure 1E), suggesting that structure in the 3′ UTR might inhibit loading of the deadenylation complex or otherwise interfere with the process of deadenylation. Meanwhile, in the presence of PABPC1, 3′ UTR structure was not observed to inhibit deadenylation (Figure S1D).

We returned to the influence that PABPC1 had on deadenylation rates and compared deadenylation rates with and without PABPC1 for each 3′ UTR tested. The addition of PABPC1 typically caused slower deadenylation rates and resulted in a narrower distribution of rates that centered at approximately 0.22 nt/min (Figure 1F). Deadenylation rates measured in the presence of PABPC1 did not correlate with those measured in its absence. Thus, PABPC1 addition appears to abrogate the effects of RNA structure that influence deadenylation when PABPC1 is not present. These observations support a model in which PABPC1 binding disrupts local RNA structure and presents the poly(A) tail to the deadenylation machinery in a more uniform and somewhat protected conformation.

### PABPC1 apparent affinities for 3′ UTRs correlate with deadenylation rates

Yeast PAB1, the ortholog of human PABPC1, contacts the 3′ UTR proximal to the poly(A) tail through RRM4^19^. Hypothesizing that human PABPC1 might similarly interact with 3′ UTRs, with varying affinity for different poly(A)-proximal sequences that could influence deadenylation rates, we measured how PABPC1 interacted with different sequences within our library through RNA bind-and-seq (RBNS)^28^. Accordingly, we incubated our library with purified PABPC1, captured bound RNA on nitrocellulose, and sequenced the enriched fraction. RNA reads from the PABPC1-bound sample were then compared to a corresponding input sample. In this analysis, tails of three different lengths were appended to the library, to simulate PABPC binding propensities at different points during deadenylation (Figure 2A). The A_30_ tail represented less extensively deadenylated RNA species, whereas the A_15_ RNA library represented more extensively deadenylated species, and the A_0_ represented fully deadenylated species.

**Figure 2.**
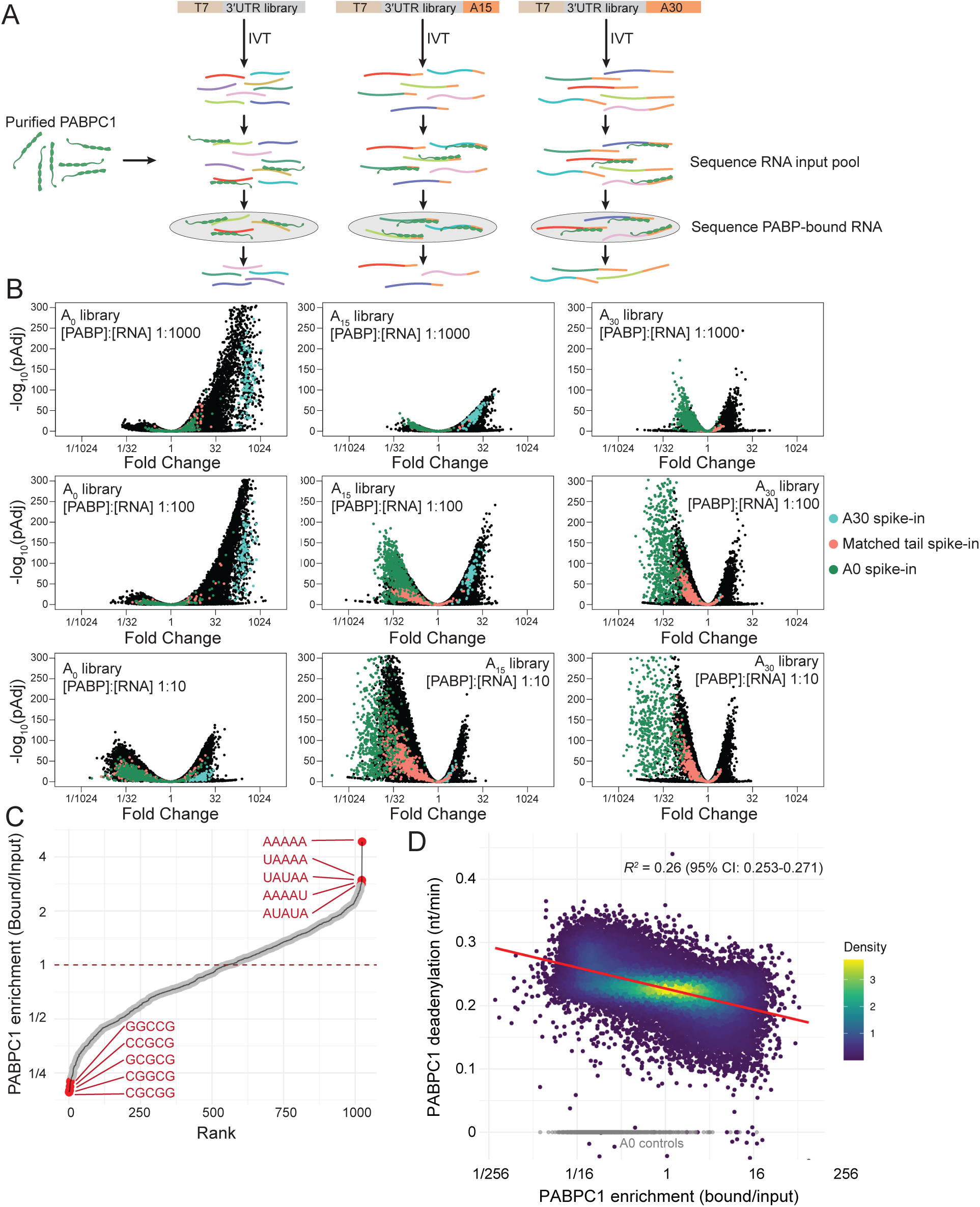
PABPC1 apparent affinities for 3′ UTRs correlate with deadenylation rates. (A) Schematic of RBNS. Three libraries of RNAs, each with a different tail length, are constructed using in vitro transcription (IVT). RNA libraries are incubated with purified PABPC1, then passed through a nitrocellulose membrane, which captures protein-bound RNA. The input library and PABPC1-bound RNA library are then sequenced. (B) Preferential enrichment and depletion of some library members. Volcano plots compare enrichment and depletion of library members in the PABPC1-bound fraction, using libraries with three different tail lengths, each assayed with three different PABPC1 concentrations. Each point represents a UTR library member, with the *p* value of its enrichment or depletion plotted as a function of its fold change in the PABPC1-bound sample compared to input. Points for RNA spike-ins with an A_30_ tail are in blue. Points for spike-ins with tail lengths matching the library tail length are in orange. Points for A_0_ spike-ins are in green. (C) Enrichment of 5-mers in library members enriched and depleted in the RBNS. Shown is the rank order of 5-mer enrichments, calculated as the ratio of reads within the PABPC1-bound fraction versus the RNA input library. The 5-mers were quantified within the A_15_ library incubated with 1 nM PABPC1 at 100-fold excess RNA library. (D) Relationship between deadenylation rate and PABPC1 enrichment in the RBNS when using the A_15_ library with 1 nM PABPC1 and a 100-fold excess of RNA library over PABPC1. 95% confidence intervals for R² were estimated by percentile bootstrap (10,000 resamples of library members with replacement).

For each RNA library with differing tail lengths, 100 nM library was incubated with limiting PABPC1, using PABPC1 concentrations of 0.1 nM, 1 nM, and 10 nM. Random-sequence control UTRs with A_0_, A_30_, and library-matched tail lengths were spiked into each experiment to calibrate expectations of PABPC1 binding (Figure S2A). After incubation for 20 h to allow binding equilibrium, RNA bound to protein was collected by using filter binding and then sequenced. As anticipated, A_30_ tail spike-in controls bound to PABPC1 with the highest apparent affinity, whereas the A_0_ spike-in controls bound with lowest apparent affinity (Figure 2B and S2D). Random-sequence spike-in controls with matched poly(A)-tail lengths bound to PABPC1 with similar but modestly lower apparent affinity than that observed for the library derived from endogenous human UTR ends. This result suggests that endogenous UTRs bind PABPC1 more effectively than randomized sequences. We noted that the randomized UTRs spanned a range of GC contents, whereas the native endogenous UTRs overall had a lower GC content than the randomized controls.

Because PABPC1 is expected to bind a 27-nt RNA footprint depending on whether the 3’ UTR sequence is present ^19^, we reasoned that differential PABPC1 binding propensities for the 3′ UTR might be difficult to observe in the A_30_ library, where direct binding to the poly(A) tail would be a dominant mode of binding. Conversely, the A_15_ library would not be long enough for all four RNA-recognition motifs within PABPC1 to bind the poly(A) tail, and thus any interactions that the 3′ UTR might make with PABPC1 would be expected to impart stronger affinity over UTRs that do not interact with PABPC1. The greatest dynamic range was observed with 1 nM PABPC1 and the A15 library. This condition was selected for further *k*-mer analysis to examine the enrichment or depletion of all possible 5-mers, which showed that A-rich and AU-containing motifs were among the motifs associated with highest PABPC1 binding propensity, whereas GC-containing motifs were enriched among sequences with the lowest binding enrichment (Figure 2C). These results identify UTR sequence features that modulate PABPC1 apparent affinity when poly(A)-tail binding becomes limiting, as occurs as a consequence of deadenylation.

We then compared the deadenylation rates in the presence of PABPC1 with PABPC1-binding enrichment measurements, revealing a negative relationship between the two (Figure 2D and S2E). Specifically, substrates with the highest PABPC1 apparent affinity tended to be deadenylated the slowest in the presence of PABPC1. These results support a model in which PABPC1 binds the poly(A) tail but can also interact with the adjacent UTR sequence, and the overall PABPC1 affinity to RNA affects the deadenylation rate.

### PABPC preferences are intrinsic to the sequence, not just to the structure of the adjacent 3′ UTR

We next explored the extent to which nucleotide sequence composition or RNA secondary structure might be influencing PABPC binding. High A content positively correlated with PABPC1 RBNS binding enrichment, whereas both GC content and predicted stability of secondary structure negatively correlated with binding enrichment (Figure 3A–C). However, because sequences with high GC content and low A content tend to form more stable predicted structures (Figure S3A,B) interpretation of these variables was confounded. To disentangle the relative contributions of nucleotide content and RNA structure, we analyzed residual correlations observed after correcting for each feature. After correcting for predicted MFE, GC content and A content each retained substantial correlation with PABPC1 apparent affinity (R^2^ = 0.17 and 0.16, respectively; Figure 3D). In contrast, after correcting for GC-content or A-content, MFE yielded much weaker residual correlations (R² = 0.03 and 0.06; Figure S3C and S3D). These results indicated that nucleotide composition was the primary determinant of PABPC1 affinity—over and above the contribution of secondary structure, consistent with PABPC1 recognizing sequence features in the 3′ UTR adjacent to the poly(A) tail.

**Figure 3.**
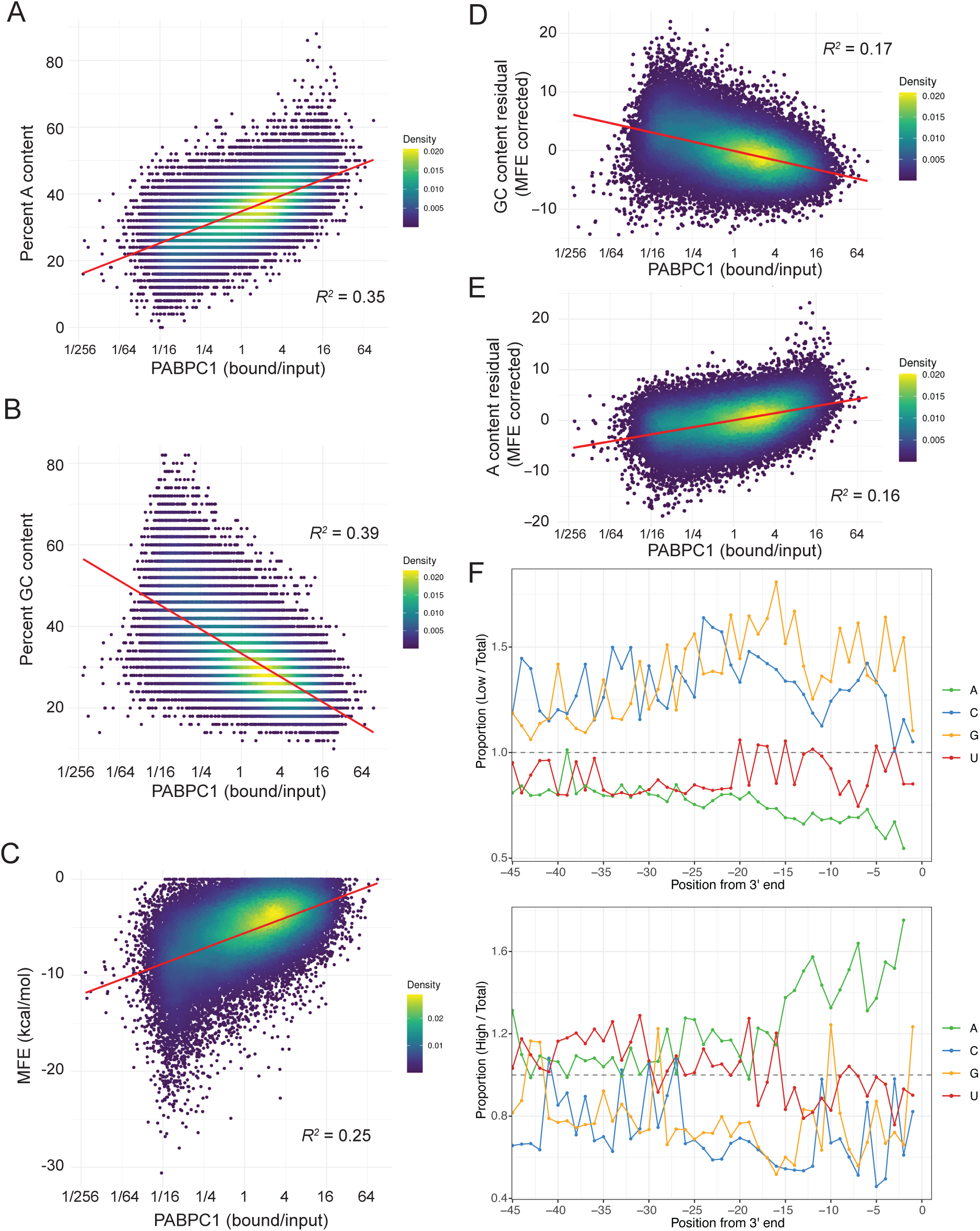
PABPC preferences are intrinsic to RNA sequence rather than RNA structure of the adjacent 3′ UTR. (A) Correlation between A content of the 50-nt UTR sequence and PABPC1 binding propensity, as measured by RBNS enrichment values for the A_15_ library at 1 nM PABPC1 and 100-fold excess RNA library. (B) Relationship between GC content of the 50-nt UTR sequence and PABPC1 apparent affinity; otherwise, as in A. (C) Relationship between predicted MFE of folding and PABPC1 apparent affinity; otherwise, as in A. (D) Residual correlation between GC content and PABPC1 apparent affinity after accounting for predicted MFE of folding; otherwise, as in B. (E) Residual correlation between A content and PABPC1 apparent affinity after accounting for predicted MFE of folding; otherwise as in A. (F) Impact of nucleotide position on PABPC1 apparent affinity. PABPC1 enrichment values (A_15_ RBNS library, 1 nM PABPC1) were divided into a low binding propensity subset (log_2_-fold change < –2.5, top) or high binding propensity subset (log_2_-fold change > 2.5, bottom). Plotted for each subset is the abundance of each nucleotide at each position in the UTR relative to the abundance of that nucleotide in the input library. Nucleotide positions were determined for each sequence by assigning position –1 to the most distal non-A nucleotide and counting backward from the 3ʹ end.

Having found that nucleotide content appeared important for PABPC1 binding preference, we investigated if the position of specific nucleotides might influence this preference. For UTRs with either low or high RBNS enrichment, the presence of a given nucleotide —A, U, G, or C—at each position was compared to the abundance of that nucleotide at that position within the input library. Poorly bound UTRs (RBNS log_2_ fold enrichment < –2.5), tended to have lower A and U content and higher G and C content throughout the UTR (Figure 3F). Well-bound UTRs (RBNS log_2_ fold enrichment > 2.5) had a strong A enrichment near the distal ends of UTRs, with the highest enrichment observed between the –15 to –2 position (Figure 3F).

### Disruption of PABPC1-UTR interaction abolishes sequence-dependent deadenylation protection

Having acquired evidence that that sequences upstream of the poly(A) tail modulate deadenylation rates through differential PABPC1 binding, we sought to confirm this model by disrupting the interaction between PABPC1 and the 3′ UTR. Structural insights^23^, combined with RRM mutation and RNA footprint mapping, suggest that RRM1 and RRM2 have a stringent selectivity for poly(A) sequence as compared to RRM3 and RRM4, which can straddle the junction of the 3′ UTR and the poly(A) tail^19^. We reasoned that replacing RRM3 and RRM4 with a second copy of RRM1 and RRM2 would disrupt UTR binding, producing a protein with strict poly(A) specificity^24^. This mutant is referred to as PABP-1212, which denotes the duplication of RRM1 and RRM2. When adding either purified PABP-1212 or purified WT PABPC1 to deadenylation reactions (Figure 4A), PABP-1212 imparted less protection against deadenylation, resulting in increased deadenylation rates (Figure 4B). Whereas addition of WT PABPC1 resulted in median rates of 0.22 nt/min (Figure 1F), addition of PABP-1212 resulted in median rates of 0.35 nt/min (Figure 4C). Although, compared to adding no PABPC, adding PABP-1212 had little effect on the median deadenylation rate (median rates of 0.37 and 0.35 nt/min, respectively), it did impart more uniform deadenylation rates (Figures 1F, 4C), presumably because it prevented the poly(A) tail from pairing with the UTR and presented the tail to the deadenylation machinery in a more uniform conformation.

**Figure 4.**
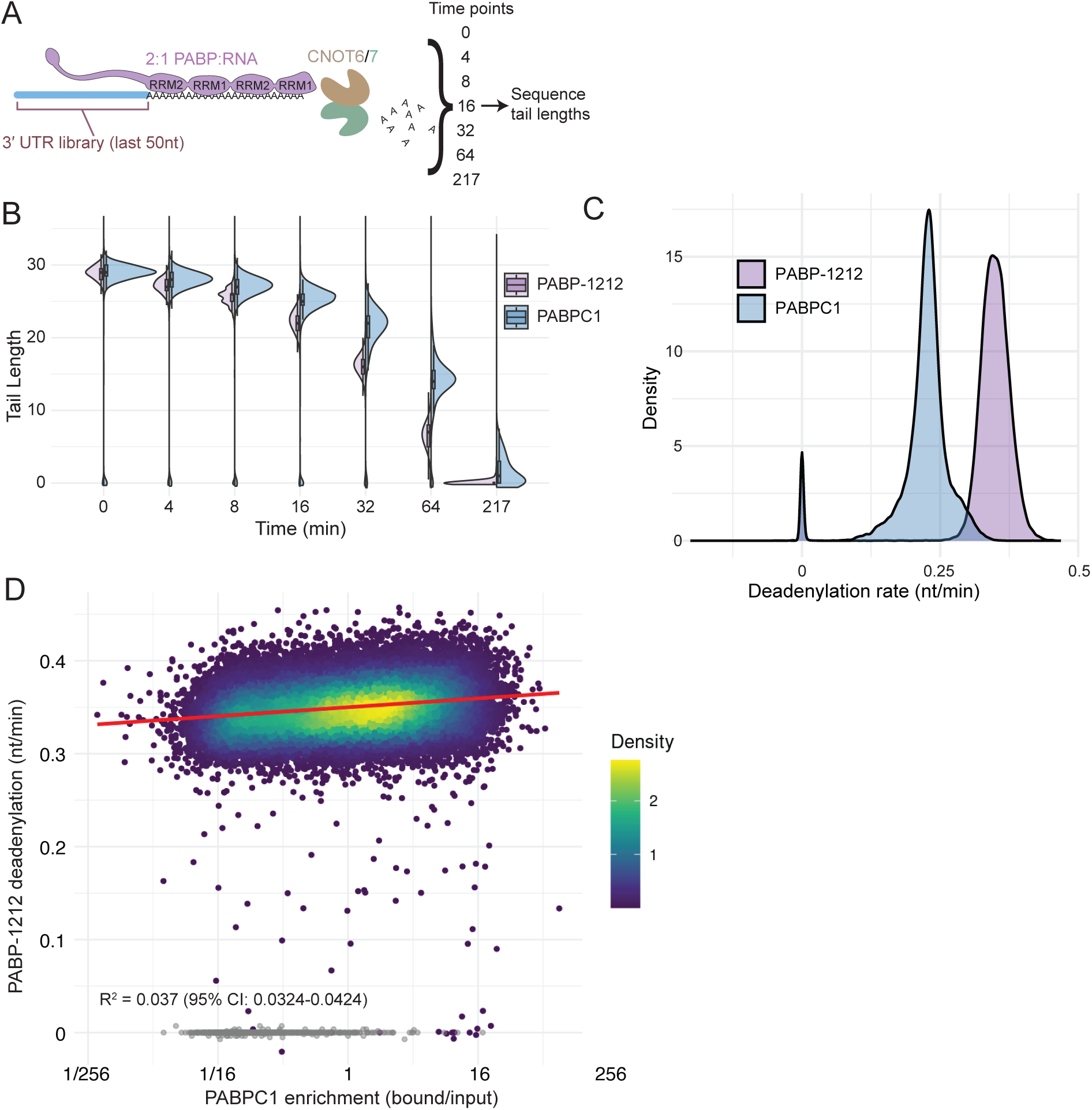
Disruption of PABPC1-UTR interaction abolishes sequence-dependent deadenylation protection. (A) Schematic of in vitro deadenylation experiment. A library of poly(A)-tailed RNA substrates is incubated with CNOT6/7 heterodimer in the presence of purified PABP-1212, a domain-substituted version of PABPC1. Otherwise, this panel is as in Figure 1A. (B) Slower deadenylation in the presence of PABPC1 versus PABP-1212. Shown for each time point are distributions of tail lengths for reactions that include either PABPC1 or PAB1212 (blue or purple, respectively). (C) Faster deadenylation in the presence of PABP-1212. Plotted are distributions of fitted deadenylation rates in the presence of PABP-1212 compared against deadenylation rates in the presence of PABPC1from Figure 1F. Peaks at zero are from the A_0_ spike-in controls within the library. (D) Poor correlation between deadenylation rate in the presence of PABP-1212 and PABPC1 enrichment in the RBNS when using the A_15_ library with 1 nM PABPC1 and a 100-fold excess of RNA library over PABPC1. 95% confidence intervals for R² were estimated by percentile bootstrap (10,000 resamples of library members with replacement).

Our model for PABPC-mediated deadenylation predicted a greater difference in deadenylation rate for WT PABPC1 vs PABP-1212 in RNA substrates preferentially bound by PABPC1. This outcome was indeed observed when comparing deadenylation rates as a function of PABPC1 apparent affinity. Compared to the clear relationship between deadenylation rate and PABPC1 binding propensity observed in the presence of PABPC1 (Figure 2D and S2E, *R*^2^ = 0.26), this relationship was nearly imperceptible in the presence of PABP-1212 (Figure 4D and S4B, *R*^2^ = 0.04). Because both proteins bind poly(A), the difference in deadenylation protection likely reflected the distinct UTR-binding properties of PABPC1 versus PABP-1212. Overall, these results examining the effects of replacing RRMs 3 and 4 with RRMs 1 and 2 further supported the conclusion that differential binding of PABPC1 RRM3 and RRM4 to the poly(A) tail modulates deadenylation rates.

### A model for PABPC1-mediated influence on deadenylation kinetics

Our data support a model in which poly(A)-proximal UTR sequence modulates the deadenylation rate through its effect on PABPC1 binding (Figure 5). Compared to UTRs with high GC content and low A content, UTRs with low GC content and high A content are more conducive to a conformation in which PABPC1 straddles the junction between the poly(A) tail and the 3′ UTR. This propensity affects the manner in which deadenylation of the last 30 nucleotides of the poly(A) tail occurs. When PABPC1 makes more favorable interactions with the 3′ UTR, it is less prone to be dislodged by the deadenylation machinery, causing deadenylation to proceed more slowly.

**Figure 5.**
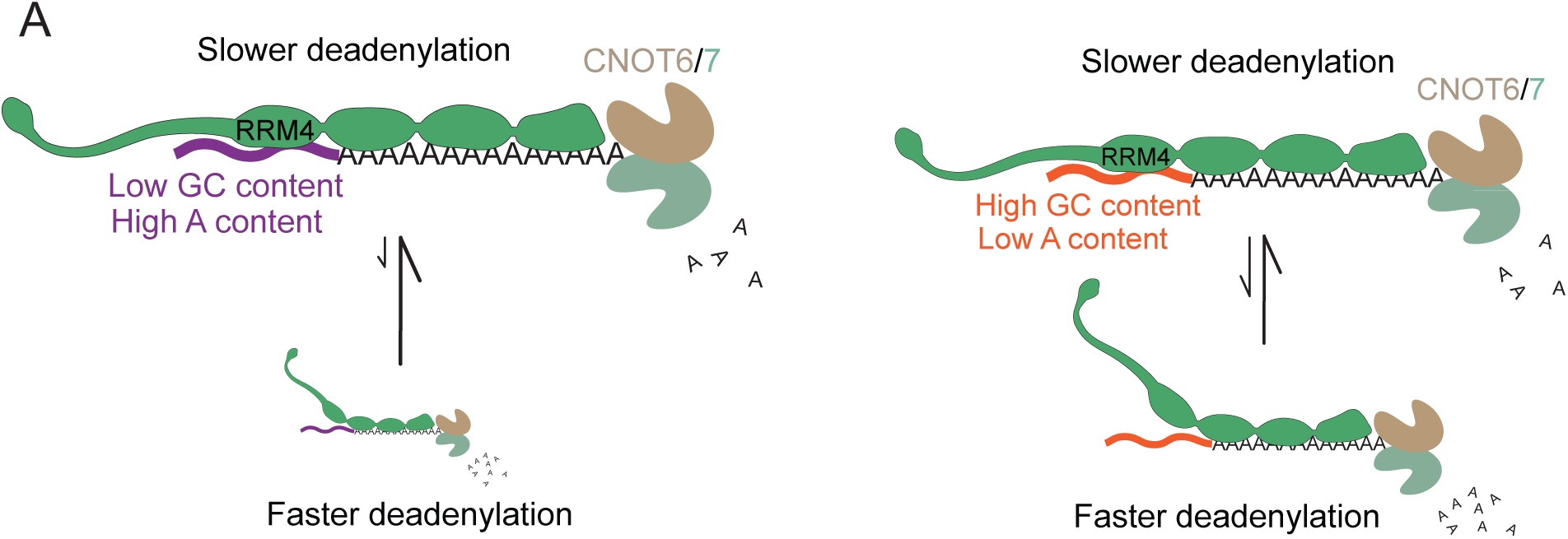
A model for PABPC1-mediated influence on deadenylation kinetics.

## Discussion

Here, we show that PABPC1 engagement with different 3′ UTR sequences contributes to transcript-specific deadenylation rates. Using a high-throughput in vitro assay, we measured deadenylation rates across thousands of human 3′ UTR sequences. RNA structure inhibited deadenylation in the absence of PABPC1, but this effect was abolished when PABPC1 was present. Even with PABPC1 present, deadenylation rates varied: PABPC1 bound more weakly to GC-rich UTR sequences, and these substrates were deadenylated faster. Domain-swap experiments confirmed that this effect is mediated by RRM3–4, which appears to enable PABPC1 to straddle the poly(A)/UTR junction. The distinction between tail-only and tail-plus-UTR binding explains how PABPC1 can preferentially stabilize specific transcripts. Together, these results show that sequences upstream of the poly(A) tail tune PABPC1 binding and deadenylation rates, likely contributing to the range of deadenylation rates observed for cellular mRNAs.

Our in vitro deadenylation assay revealed that RNA structure near the poly(A) tail can inhibit deadenylation in the absence of PABPC1, but that this inhibition is relieved by PABPC1 binding. We speculate that PABPC1 engages these nucleotides, overriding local RNA structure and presenting the poly(A) tail in a more homogeneous configuration for deadenylation. Consistent with this model, the distribution of deadenylation rates without PABPC1 showed a peak at ∼0.45 nt/min corresponding to unstructured substrates, with a broad shoulder toward slower rates that correlated with increasing predicted MFE. PABPC1 collapsed this distribution into a single peak at ∼0.22 nt/min. Such a model would suggest that structure-dependent regulation of decay might often require limiting PABPC activity.

PABPC1 abundance varies across cell types and conditions, which may allow other determinants of mRNA fate, including structure-dependent regulation, to exert their effects. In post-embryonic cells, PABPC1 is estimated to be at 7:1 molar excess compared to poly(A) tails, or 2:1 molar excess compared to number of total sites within the poly(A) tail^29^. The concentration of PABPC1 is also approximately 1000-fold above its measured K_d_ of 4 nM^30,31^, so poly(A) tails in these cells are expected to be fully occupied by PABPC1. However, in oocytes and early embryos, and in coronavirus-infected cells, PABPC is limiting—particularly for short-tailed mRNA isoforms^32–34^—creating conditions in which 3′-UTR structure would be able to play a role in modulating deadenylation. Consistent with this model, PABPC abundance is reported to tune the susceptibility of mRNAs to RBP-directed deadenylation in a concentration-dependent manner^35^, suggesting a context of limiting PABPC that might similarly permit structure-dependent regulation.

Even in the more typical regime, in which PABPC is saturating poly(A) tails, we observed differential deadenylation kinetics that corresponded to RBNS-determined differential PABPC1-binding propensities. Perhaps faster PABPC dissociation rates at lower-affinity UTRs enable more frequent, albeit transient, access to the deadenylation machinery, even if PABPC1 is usually occupying the poly(A) tail. Despite low K_d_ and apparent full occupancy, variation in k_off_ might produce measurable differences in deadenylation rates.

These findings have several implications for gene regulation. First, PABPC1 is implicated as a transcript-specific regulator of mRNA decay—not simply a global factor. Second, these findings help explain the mechanisms that give rise to the wide range of deadenylation kinetics across transcripts. Third, the ability of 3′ UTRs to encode PABPC1-dependent protection suggests that selective stabilization could contribute to coordinated expression of gene programs. Two distinct mechanisms enable selective stabilization, depending on whether PABPC1 is limiting. These findings may extend to other PABPC1-dependent processes, such as its role in translation regulation. For example, PABPC1 acts during early embryogenesis, where it can influence translation efficiency and is also an important host factor in the retrotransposition of LINE-1 elements^16,36^.

The sequence-dependent effects we observe explain only a modest fraction of the variation in cellular deadenylation rates. The effects described here are ∼1.3-fold in magnitude, whereas cellular decay rates span a nearly 1000-fold range. Thus, poly(A)-proximal sequence represents one contributor among many that collectively shape mRNA stability. In addition, the influence of UTR-encoded PABPC1 apparent affinity is likely most pronounced when poly(A) tails are short, a state that constitutes only a subset of mRNA molecules at any given time for most genes. This mechanism may therefore operate within a defined temporal window of the mRNA lifecycle, contributing alongside other cis-elements, trans-acting factors, and cellular conditions to the full range of decay kinetics observed in vivo.

How do interactions of PABPC1 with other decay or translation factors intersect with its sequence-specific binding at the UTR–tail junction? Do variant residues of PABPC1 paralogs or post-translational modifications alter these sequence preferences? Are fluctuations in PABPC abundance across embryogenesis or during T-cell activation^32,37^, harnessed to regulate specific transcripts? Addressing these questions will clarify the scope of PABPC1-UTR interactions in gene regulation.

## Methods

### Plasmid construction

Plasmid pRM010c was cloned for PABPC1 protein over-expression in *E. coli*. Full-length human PABPC1 coding sequence was codon-optimized for E. coli expression and synthesized as three gene blocks with overlapping homology (IDT). These gene blocks were assembled by Gibson reaction into a pET28a vector encoding PABPC1 with an N-terminal His6-SUMO tag for affinity purification and tag removal by Ulp1 protease.

Plasmid pRM039 was similarly cloned for PABP-1212 protein over-expression in *E.coli*. PABP-1212 was designed as a PABPC1 domain substitution in which the region coding for RRM3, the intervening protein linker, and RRM4 of a *E.coli* codon-optimized PABPC1 was replaced with RRM1, the intervening protein linker, and RRM2 of *E.coli* codon-optimized PABPC1. Three segments of the PABP-1212 sequence were amplified by PCR with homology arms and assembled by Gibson reaction into the pET28a vector with tandem N-terminal HIS tag + SUMO tag.

### Purification of PABPC1 and PABP-1212

*E. coli* strain BL21-DE3 Rosetta2 was transformed with either plasmid pRM010c or pRM039, and each transformation was plated on agar containing both chloramphenicol and kanamycin. Multiple colonies were taken from each plate, pooled together, and used to seed 100 mL liquid cultures which were incubated overnight at 30°C with 225 rpm shaking. These liquid cultures were then used to seed four 1.5 L liquid cultures in baffle flasks at 1/100 dilution, which were incubated at 37°C with 225 rpm shaking. When cultures reached OD = 0.2, they were transferred to 16°C with 225 rpm shaking. When these chilled cultures reached OD = 0.6, they were induced with 0.5 mM IPTG, grown for an additional 20 h at 16°C with 225 rpm shaking, and harvested by pelleting via centrifugation at 5000*g* for 15 min. Pellet volume was estimated by weight and two volumes of lysis buffer were added on ice. Lysis buffer contained 50 mM Tris pH 8.0, 2 M NaCl, 5% glycerol, 0.5% NP-40, 15 mM imidazole, 4 mM BME, and one tablet of Roche mini protease inhibitor cocktail added for every 100 mL buffer. The high NaCl concentration inhibited downstream co-purification of RNA with PABPC. Bacterial protease-inhibitor cocktail (P8465-5ML) was prepared from lyophilized powder by adding 1 mL DMSO and 4 mL water, then added at 1 mL per 20 mL of cell lysate. This lysate was sonicated at 50% on/off with 1 s pulses for 0.5–1 min per round, with 1 min rest on ice so as not to overheat the sample. Sonication was performed for five rounds until the lysate became free-flowing. Lysate was clarified with 40,000*g* centrifugation at 4°C for 30 min, and the supernatant was transferred to a separate tube.

Nickel-agarose beads (1 mL beads for every 8 L bacterial liquid culture) were pre-equilibrated with lysis buffer by spinning at 100*g* at 4°C, removing supernatant, and resuspending in lysis buffer. These pre-equilibrated beads were then added to the clarified lysate and rotated end-over-end at 4°C for 1 h. Bead slurry was applied to a gravity flow column, allowed to drain, and the beads washed three times with lysis buffer. Protein was then eluted using 3 mL elution buffer (50 mM Tris pH 8.0, 500 mM NaCl, 5% glycerol, 0.5% NP-40, 250 mM imidazole, 4 mM BME). Beads and column were saved for rebinding of HIS-SUMO tandem tag. Purified ULP1 protease was added to the eluant and this mixture was dialyzed overnight at 4°C in dialysis buffer (50 mM Tris pH 8.0, 200 mM NaCl, 5% glycerol, 0.5% NP-40, 15 mM imidazole, 4 mM BME). Ni beads were equilibrated with two washes of dialysis buffer and the dialyzed eluant was reapplied to capture cleaved HIS-SUMO tandem tag. Flowthrough was collected, together with an additional 1 mLwash of dialysis buffer. Protein was further purified by ion exchange on a heparin column (low-salt buffer: 20 mM Tris pH 8.0, 200 mM NaCl, 5% glycerol, 5 mM DTT; high-salt buffer: 20 mM Tris pH 8.0, 2 M NaCl, 5% glycerol, 5 mM DTT). Protein was then concentrated by size exclusion on a Superdex 200 column. Coomassie stains were used throughout the purification, for diagnostics and to identify fractions containing full-length protein.

### Design and preparation of RNA libraries

The collection of human poly(A) sites was assembled from annotations in polyA_DB version 3.2, which catalogs poly(A) sites that have been observed in RNA-seq datasets. This collection was chosen on the basis of conservation (as those conserved in at least two other species, among mouse, rat, and chicken) and expression (the 38,700 sites with the highest mean RPM (average reads per million of PAS reads across all human samples included in the database). A DNA library corresponding to the 50-nt sequence segments upstream of the poly(A) tail in these 38,700 sites with three different tail lengths (A_0_, A_15_, and A_30_) was synthesized by Agilent. Each tail length was designed with unique PCR flanking sequences to allow for separate preparation of each tail length RNA substrate. A_0_ substrate used 5′ flanking sequence CTACACGACGCTCTTCCGATCTCCTGAG and 3′ flanking sequence GGCCGGCATGGTCCCAGCCTCCTCGCTGGC. A_15_ substrate used 5′ flanking sequence CTACACGACGCTCTTCCGATCTGGACTC and 3′ flanking sequence GCCCGGGATGGTCCCAGCCTCCTCGCTGGC. A_30_ substrate used 5′ flanking sequence CTACACGACGCTCTTCCGATCTGTGCCA and 3′ flanking sequence GCGGCCGATGGTCCCAGCCTCCTCGCTGGC.

Three separate PCR reactions were performed on the synthesized library, each using a different set of oligos that recognized each unique 5′ and 3′ flanking sequence. Each PCR reaction also added filler sequence, the T7 promoter, and TruSeq R1 sequence on the 5′ end and an HDV ribozyme sequence plus filler sequence at the 3′ end. Each PCR product was amplified for 16 cycles with NEBNext Ultra II Q5 DNA polymerase, and purified on a 3% MetaPhor agarose gel. Purified DNA was used as a template for in vitro transcription and heat cycled to allow for efficient HDV ribozyme cleavage that removed sequence beyond the end of the encoded poly(A) tail, resulting in RNA substrates containing a 5′ TruSeq R1 sequence, a 50nt sequence corresponding to one of the poly(A) proximal sequences within the library, and an A_0_, A_15_, or A_30_ tail. These RNAs were purified on a denaturing polyacrylamide gel and treated with T4 polynucleotide kinase to remove the 2′-3′ cyclic phosphate created during HDV ribozyme cleavage. RNA substrates were then purified by phenol–chloroform extraction and used for downstream experiments.

### Deadenylation assay

The A_30_ RNA substrate library (200 nM) was pre-incubated at 37°C for 20 h with either PABPC1 (400 nM), PABP-1212 (400 nM), or no PABPC1 in binding buffer (16 mM HEPES pH 7.4, 89 mM potassium acetate, 0.89 mM magnesium acetate, 0.87 mM Tris HCL pH 7.5, 1.7 mM NaCL, 0.0029% IGEPAL-CA-630, 0.0089 mg/mL yeast tRNA, 0.029 mg/mL BSA, 7mM DTT, 1 U/µL SUPERase-in, and 8% v./v. glycerol). Purified CNOT6/7 heterodimer^38^ was diluted into the same binding buffer and added to the pre-incubated PABP-RNA substrate library, resulting in a final concentration of 100 nM RNA substrate, 200 nM PABPC, and 200 nM CNOT6/7 heterodimer. Time points were collected and quenched with five volumes of 4°C PCA. RNA was then extracted and sequenced to measure tail lengths for individual 50-nt poly(A)–proximal sequences.

### PABPC1 RBNS

The A_0_, A_15_, and A_30_ RNA libraries (100 nM final concentration) were each incubated with three concentrations of PABPC1 (0.1 nM, 1 nM, and 10 nM final concentrations) for 20 h at 37°C, in binding buffer used for deadenylation assays. In a 37°C warm room, 10 µL of each binding reaction was then applied to a filter-binding suction cassette, comprised of a circular nitrocellulose filter punch-out placed on top of a Hybond circular punch-out. In this double-filter set-up, the nitrocellulose membrane captured PABPC1-bound RNA, and the Hybond nylon membrane captured any RNA that passed through the nitrocellulose filter. After a single wash with 100µL of ice-cold wash buffer (18mM HEPES pH 7.4, 100mM potassium acetate, 1mM magnesium acetate) under suction, the nitrocellulose membranes were collected. Each membrane was placed in a 1.5 mL RNAse-free G-tube, and 420 µL proteinase K reaction mix (50 mM Tris-HCl pH 7.5, 50 mM NaCl, 10 mM EDTA, 1% SDS, 1µg/µL proteinase K, 0.01 mg/mL yeast tRNA) was added to each tube. Reactions were incubated in a thermomixer at 65°C for 45 min with 1200 rpm shaking. RNA from the reactions was then phenol–chloroform extracted, ethanol precipitated, and processed for high-throughput sequencing. RNA was also phenol–chloroform extracted and ethanol precipitated from 10 µL of the RBNS input binding reaction, and processed for sequencing in parallel to the filtered samples.

### Sequencing of RNA libraries

Pre-adenylated oligo adapter was ligated to the 3′ end of extracted RNA using T4 RNA ligase 1 (NEB) and incubating at 23°C for 150 min. 20 pmol of pre-adenylated oligo adapter was sufficient to ligate RNA extracted from either 10 µL of RBNS input, one RBNS nitrocellulose membrane, or 10 µL of quenched deadenylation reaction. After phenol-chloroform extraction and ethanol precipitation, RNA was annealed to primer GGCTCGGAGATGTGTATAAG and reverse transcribed using SuperScript III (Invitrogen). The RNA–cDNA duplex was purified and concentrated using DNA Clean (Zymo) with a 7:1 ratio of binding buffer, which removed free primer and primer-free adapter duplexes. This cDNA was then used as the input for PCR amplification with NEBNext Ultra II Q5 DNA polymerase and dual-indexed primers that recognize the TruSeq binding site on the forward strand and the Nextera binding site on the reverse strand. For each reaction, different cycle numbers were tested, ranging from 12 to 22. These reactions were resolved on a MetaPhor agarose gel, and the band at the expected size was extracted for the lowest cycle number that produced a visible single band without also producing a higher band diagnostic of PCR over-amplification. Amplicons from RBNS experiments were submitted for high-throughput 100-nt single-end sequencing on the NovaSeqS1 (Illumina). Amplicons from deadenylation experiments were submitted for high-throughput 300-nt single-end sequencing on the AVITI (Element). Both sequencing datasets generated during this study will be available at GEO [Accession Number Pending].

### Deadenylation rate estimation

Deadenylation samples were de-multiplexed using dual indexed AVITI sequencing. Flanking constant sequence was trimmed from each read using Cutadapt^39^ to extract the 50-nt sequence adjoining the poly(A) tail. Due to increased sequencing error that might occur on the poly(A) homopolymer, the end of the poly(A) tail was defined as the position at which two or more contiguous non-A nucleotides were detected. Reads were then aligned to the reference file of 50-nt poly(A)-proximal sequences using Bowtie2^40^, and a custom python script was used to calculate poly(A)-tail length from the unaligned 3′ end of the read. For each library member, the median poly(A) tail length was calculated across all reads mapping to that member at each timepoint. Deadenylation rates were estimated by ordinary least-squares linear regression of median poly(A) tail length against time, using six timepoints (0, 4, 8, 16, 32, and 64 min). The 217-min timepoint was excluded from rate estimation because the majority of substrates had been fully deadenylated by this time, creating a floor effect that would bias linear rate estimates. Data from both biological replicates were included in each regression (n = 12 observations per library member per PABP condition: 6 timepoints × 2 replicates), and the resulting slope was taken as the deadenylation rate (nt/min). Library members were retained for downstream analysis only if the combined read count across both replicates at the initial timepoint (t = 0) exceeded 10 for each PABP condition (WT, noPABP, and PABP-1212). Library members with missing data at any timepoint were excluded. For comparisons of deadenylation against RNA structure, RNAfold was used to calculate MFE^27^.

### RBNS enrichment analysis

RBNS samples were de-multiplexed using dual indexed AVITI sequencing. Flanking constant sequence was trimmed from each read using Cutadapt^39^ to extract the 50-nt poly(A)-proximal sequence. Bowtie2^40^ was used to align these reads to the reference set of library members and count abundance within each sample. Differential enrichment of library members in the PABPC1-bound (nitrocellulose filter-retained) fraction relative to input was quantified using DESeq2^41^. For each combination of poly(A) tail length (A0, A15, A30) and PABPC1 concentration (0.1, 1, 10 nM), read counts for each library member across input and filter-bound fractions were modeled separately using the design formula ∼ sample_type, where sample type denotes input or filter-bound. Two biological replicates were performed for each condition. DESeq2’s default median-of-ratios method was used for library size normalization, and dispersion estimates were obtained using DESeq2’s empirical Bayes shrinkage. P-values were adjusted for multiple testing using the Benjamini-Hochberg procedure as implemented in DESeq2, and DESeq2’s automatic independent filtering was applied to remove low-count library members with insufficient statistical power prior to p-value adjustment. Log₂ fold-change values from these comparisons are referred to as RBNS enrichment scores throughout.

## Acknowledgements

We thank the Whitehead Institute Genome Technology Core for high-throughput sequencing. R.Y.M. was a Howard Hughes Medical Institute fellow of the Damon Runyon Cancer Research Foundation (grant no. DRG-2485-23). D.P.B. is an investigator of the Howard Hughes Medical Institute. This study was supported by the Intramural Research Program of the National Institutes of Health (project number 1ZIABC011977 to E.V.) The contributions of the NIH authors are considered works of the United States Government. The findings and conclusions presented in this paper are those of the authors and do not necessarily reflect the views of the NIH or the U.S. Department of Health and Human Services. We thank J. Lobel for fruitful discussions and providing feedback on the manuscript. We thank A. Latifkar, B. Wierbowski, and other members of the Bartel lab for fruitful discussions.

---

**Figure S1.**
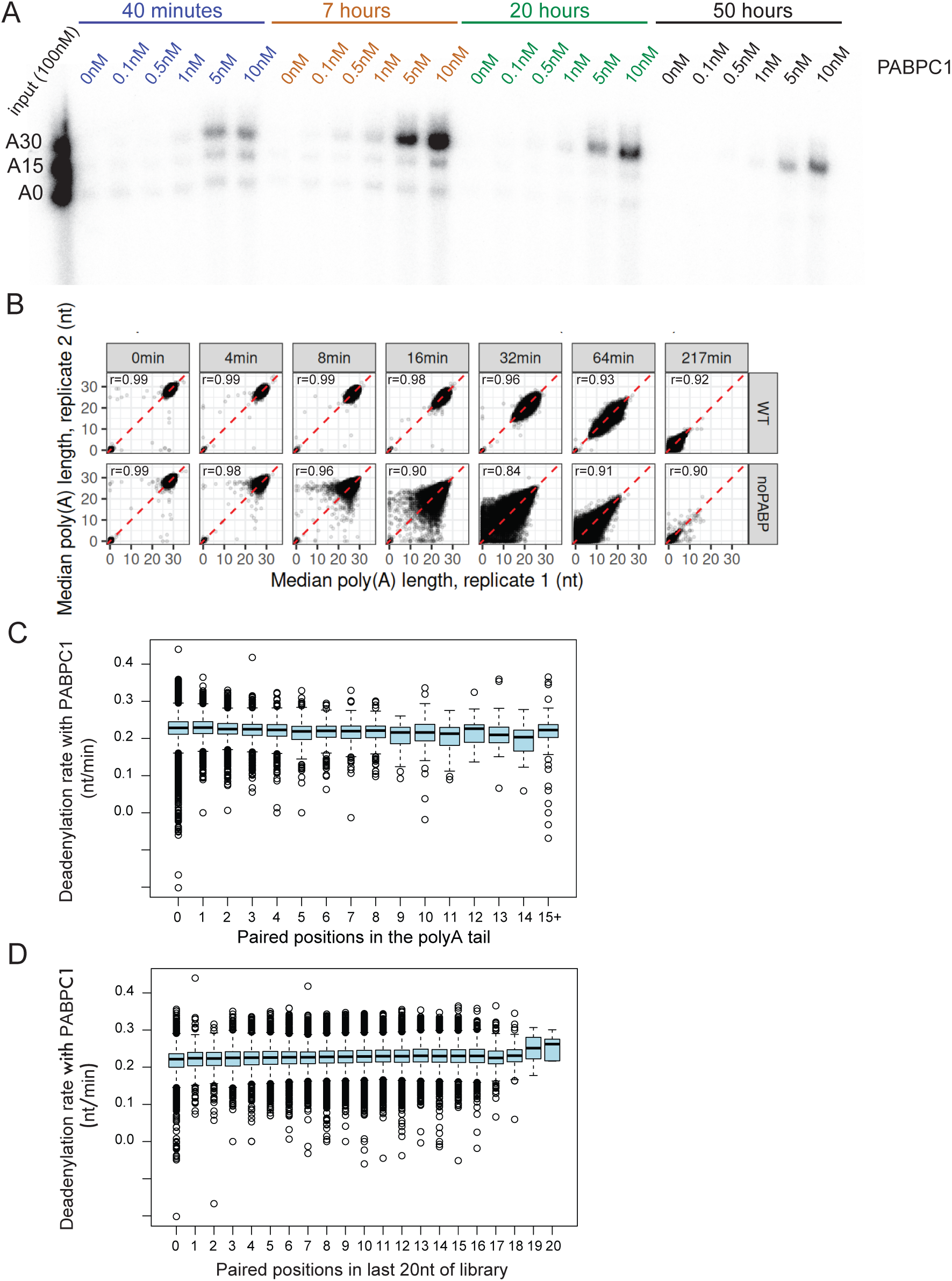
In the presence of PABPC1, UTR structure does not appear to affect deadenylation in vitro. Related to **Figure 1**. (A) Time required for PABPC1 to approach binding equilibrium. 100 nM even-ratio mix of A0, A15, and A30 radiolabeled RNA substrate was incubated with titrated levels of PABPC1 for various lengths of time. Filter binding was used to assess RNA binding to PABPC1. (B) Replicate plots of median poly(A) tail length across deadenylation time points in the presence (top) or absence (bottom) of PABPC1. (C) In the presence of PABPC1, negligible change in deadenylation rates as tail residues predicted to be paired increase. Box and whisker plots show the distributions of fitted deadenylation rates in the presence of PABPC1 for library molecules with the indicated number of poly(A)-tail positions that are predicted to be paired in the lowest-energy structure (line, median; box quartile; whisker, 1.5-times IQR rule via Tukey’s method). (D) In the presence of PABPC1, negligible change in deadenylation rates of molecules with more tail-proximal residues that are predicted to be paired. This panel is as in A, but instead groups deadenylation rates by the number of positions within 20 nt preceding the poly(A) tail that are predicted to be paired in the lowest-energy structure.

**Figure S2.**
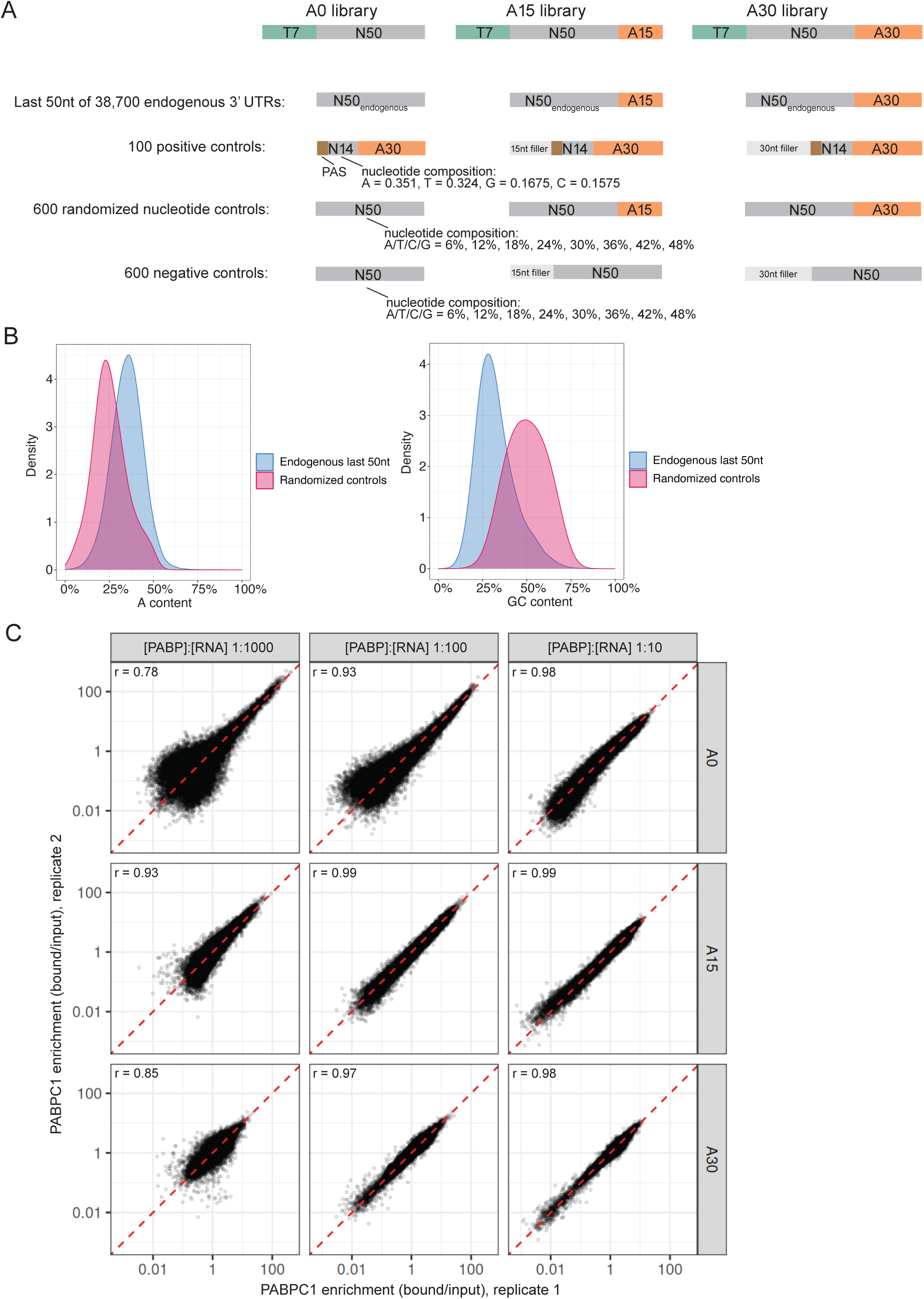
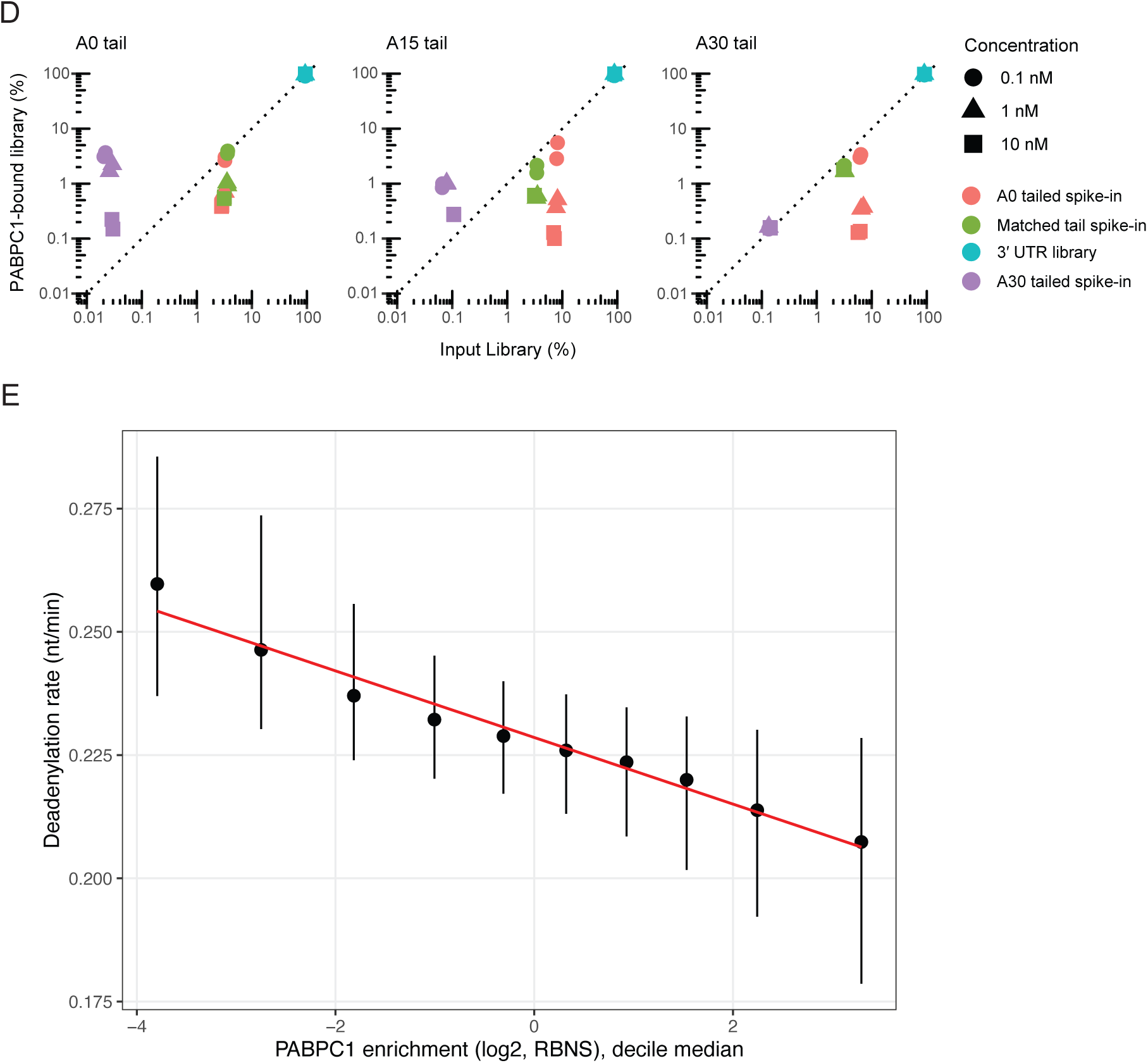
RNA substrate spike-in design and bulk PABPC1 binding propensity properties. Related to **Figure 2**. (A) RNA library design with spike-in controls. Positive controls contained an A_30_ tail, and negative controls lacked a poly(A) tail. N50_endogenous_ represent human UTR sequences, whereas N14 and N50 regions are random-sequence regions that were then filtered to a set that achieved the specified nucleotide composition. N14 sequences had the same nucleotide composition as human 3′ UTRs, whereas N50 sequences had a spread of nucleotide compositions such that at least 20 sequences represented each nucleotide at each of the specified percentages (6, 12, 18, 24, 30, 36, 42, 48). (B) The distribution of A-content and GC-content across the designed library and randomized control sequences. (C) Replicate plots of PABPC1 enrichment (bound/input) from RBNS across tail lengths and PABPC1 concentrations. (D) Enrichment plots comparing the input library against the PABPC1-bound library. PABPC1 concentration titration is depicted as circle, triangle, or square (key) and sample descriptors are depicted with orange, green, blue, or purple color. (E) Magnitude of effect visualized by quantile-binned comparison of deadenylation rate in the presence of PABPC1 as a function of PABPC1 RBNS enrichment.

**Figure S3.**
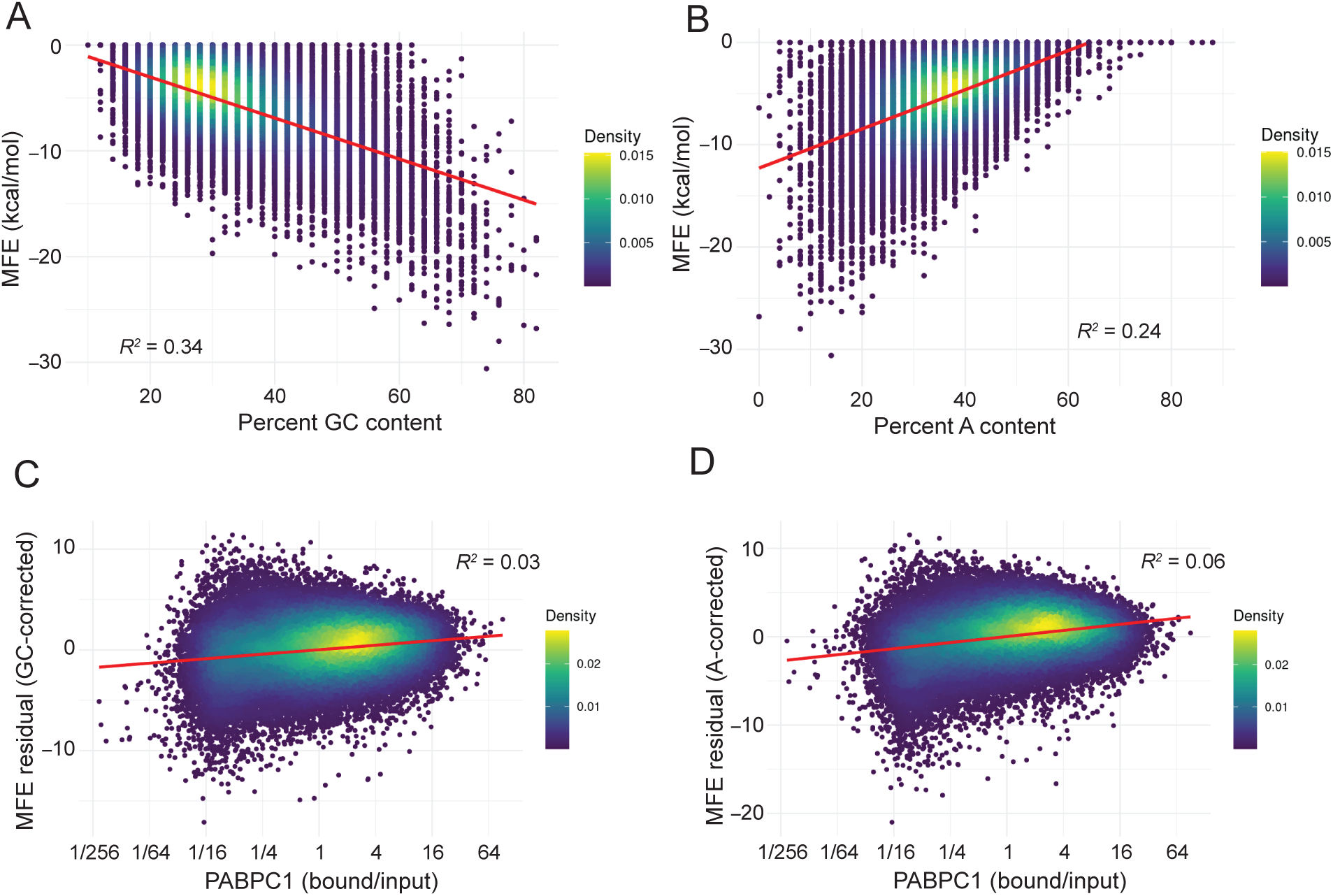
MFE residual after correction for nucleotide content shows little correlation with PABPC1 apparent affinity. Related to **Figure 3**. (A) Negative correlation between MFE (calculated for each 50 nt UTR with adjoining A_15_ tail) and percent GC content. Each point in the scatterplot represents a member of the RNA library. (B) Positive correlation between MFE and percent A content; otherwise in A. (C) Negligible correlation between MFE with GC-content correction and PABPC1 apparent affinity, measured as enrichment in the PABPC1-bound fraction (A_15_ library at 1 nM PABPC1). (D) Poor correlation between MFE with A-content correction and PABPC1 apparent affinity; otherwise, as C.

**Figure S4.**
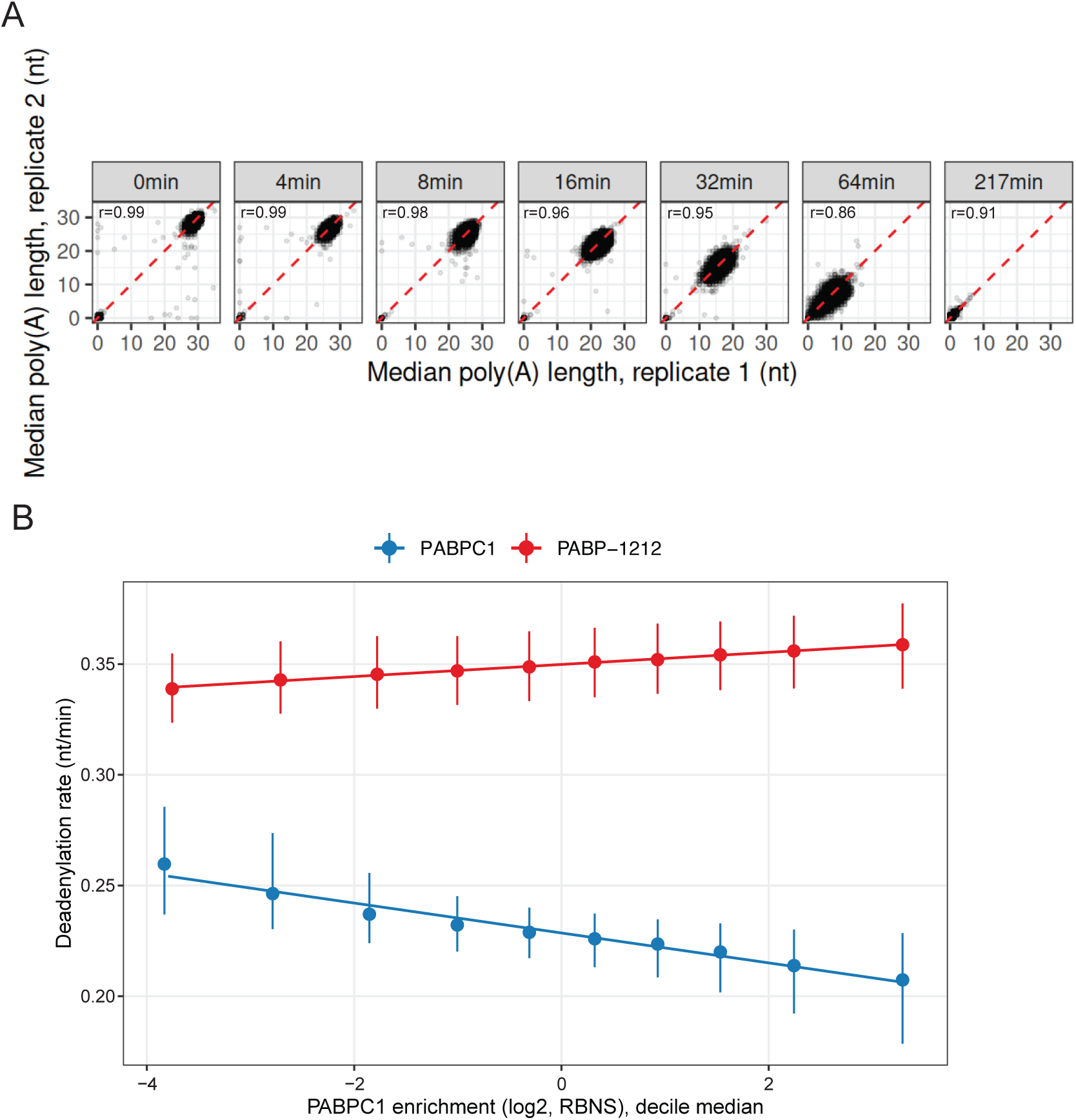
A PABPC1 domain-swap mutant that loses UTR binding also loses sequence-associated deadenylation protection. Related to **Figure 4**. (A) Replicate plots of median poly(A) tail length across deadenylation time points in the presence of PABP-1212. (B) Magnitude of effect visualized by quantile-binned comparison of deadenylation rate in the presence of PABP-1212, compared to PABPC1, as a function of PABPC1 RBNS enrichment.

## Notes

### Competing Interest Statement

The authors have declared no competing interest.

